# Development of Non-invasive ABPP-BRET Platform Technology Enable Real-Time Detection of Post-Translational Modification and Target Engagement with Absolute Specificity in Living Systems

**DOI:** 10.1101/2022.05.26.493547

**Authors:** Punita Bathla, Aaiyas Mujawar, Abhijit De, Britto S Sandanaraj

## Abstract

Non-invasive, real-time, longitudinal imaging of protein functions in living systems with unprecedented specificity is one of the critical challenges of modern biomedical research. Despite several advancements, it is estimated that nearly 35% of the human proteome is not completely characterized. Therefore, the development of new technologies is imperative for shining more light on so-called “dark proteomes”. Towards that goal, here we report a platform fusion technology called activity-based protein profiling-bioluminescence resonance energy transfer (ABPP-BRET). This method provides an opportunity to study the post-translational modification of a target protein in real-time in living systems in a longitudinal manner with a high spatio-temporal resolution. This semi-synthetic BRET biosensor method is used for target engagement studies and further for inhibitor profiling in live cells. The simplicity of this method coupled with the critical physical distance dependent BRET read-out turned out to be a powerful method, thus pushing the activity-based protein profiling technology to the next level.

## Introduction

Proteins play a major role in both human physiology and pathology. Despite their critical role, nearly 35% of the human proteome are still not characterized. Out of 65% of proteomes that are characterized, less than 5% of the proteome is focused on drug discovery programs.^1^ However, there is renewed interest to study the entire human proteome in an unbiased manner. The development of new proteomic technologies would greatly facilitate annotating the function of a plethora of proteins. In this regard, chemical proteomic technologies have gained a lot of attention in the last two decades.^2,3^ In particular, the activity-based protein profiling (ABPP) method has revolutionized the way that we study protein function^4,5^. Currently, this method is widely used for assigning new functions for both known and unknown proteins^4,5^. In addition this method is widely used in pharmaceuticals/biotech industries and academic laboratories for target identification, target validation, and inhibitor screening^6–12^. ABPP method uses gel- and/or mass spectrometry (MS)-based detection platforms. The use of gel-based detection is highly attractive because of its simplicity and it is an inexpensive setup. However, the ABPP-Gel method suffers from poor resolution and narrow dynamic range. On the other hand, the MS-based detection method offers superior sensitivity and provides the ability to profile multiple proteins simultaneously in an unbiased manner. Despite these attractive features, the ABPP-MS method suffers from extremely low-throughput which limits parallel and reproducible sample analyses, laborious sample preparation, and the need for large input proteome. Further, the ABPP-MS method requires homogenization of cells/tissues, which results in the loss of spatial information about protein activity both at intra- and inter-cellular levels.

To address some of the limitations of ABPP-MS, others and we have developed complementary technologies. Moellering group introduced a new method named the “activity-dependent proximity ligation (ADPL)” platform^13,14^. ADPL integrates the use of activity-based probes along with barcoded oligonucleotide proximity ligation and amplification. ADPL offers several advantages such as (i) direct visualization of “active proteins” with a high spatial resolution, (ii) increased dynamic range through signal amplification, and (iii) deciphering phenotypic heterogeneity in patient tissues. Further, they extended this method for profiling many enzymes simultaneously through multiplexing^15^. The multiplexed activity-based approach was applied to directly quantify small molecule-protein target engagement^15^. Although, this method addresses some of the limitations of ABPP-MS, it has several drawbacks such as (i) it requires fixing and permeabilization of cells followed by treatment with antibodies labeled with oligonucleotides; (ii) requires the availability of high-quality primary antibodies that can bind to target proteins with high affinity and specificity; (iii) it is not guaranteed that the epitope region of target protein always be available for antibody recognition as it may be occupied by other biomolecules in the highly complex environment of a living cell; (iv) this method also requires preparation of secondary antibodies labeled with oligonucleotides making it more laborious. Most importantly, and (v) this method cannot be used to study the function of active proteins in living systems in real-time in a longitudinal manner.

During the same time, our group was also working on the development of complementary technologies to ABPP-MS. We developed a powerful yet simple and easy method named “activity-based reporter gene technology (AbRGT)”^16,17^. As the name suggests this method is the fusion of ABPP with the reporter gene technology. Our technology uses fluorescence resonance energy transfer (FRET) measurement for the detection of protein activity. Using this technology, we demonstrated imaging of active protease in a single live cell in real-time with unprecedented specificity. We have used this method to study the function of different proteases such as caspase-3, - 7, -8, and -9 in an apoptosis signaling pathway. In addition, for the first time, we investigated the role of cathepsin-B in an apoptosis pathway in a non-invasive manner^16^. Further, this method was used for target engagement studies and inhibitor profiling in live cells. Compared to ABPP-MS and ADPL technologies, ABRGT offers several advantages such as (i) extremely simple and powerful method, (ii) offers protein profiling in live cells with unprecedented specificity, (iii) real-time imaging capabilities, (iv) longitudinal imaging, and (v) in principle could be adopted to study the function of any protein (platform technology). Although extremely powerful, like any other method, this technology also has a few limitations such as (i) the need for highly sophisticated instruments such as a confocal microscope, (ii) narrow dynamic range, (iii) low-throughput, (iv) the need for highly trained personnel and (v) most importantly this technology may not be translated to protein profiling in live animal as FRET study may not be ideal for whole-body imaging experiments.

To address the limitation of our previous method, here we report a platform technology, named ABPP-BRET. We demonstrate the capability of this method to follow post-translational modification of the caspase-3 enzyme in live cells in real-time in a longitudinal way. This method was used for real-time target engagement/inhibitor profiling in live cells by creating a stable cell line expressing caspase-3-luciferase fusion protein and followed by monitoring of caspase-3 activation upon treatment with a pan-kinase inhibitor. Most importantly, this technology provides opportunities to translate ABPP to study protein function with an unprecedented specificity.

The ABPP-BRET method is schematically depicted in Figure 1. Here, the protease-of interest (PoI) is cloned into a plasmid vector for mammalian expression encoding the luciferase gene generating a luciferase-tagged PoI recombinant plasmid (Figure 1a). Luciferase-tagged PoI plasmid is expressed in a suitable cell line (Figure 1b.). An appropriate stimulus is applied to the cells for the activation of PoI (Figure 1c). Cells were then incubated with an activity-based fluorescent probe (ABFP) for the labeling of PoI (Figure 1d). The choice of ABFP should be such that it allows a significant spectral overlap between its excitation and the emission of luciferase. Subsequently, specific luciferase substrate is added to the cells. The luciferin substrate will be acted by luciferase leading to the generation of photons. As the excitation of ABFP overlaps with the emission spectra of the luciferase the ABFP gets excited, resulting in the BRET effect (Figure 1e). Since the biophysical principle of BRET process is proximity dependent, the efficiency of transfer of energy is dependent on physical distance between the donor (excited luciferase substrate) and acceptor (ABFP). The non-specific labeling of other proteases by ABFP in the cell will not interfere with the BRET signal. Thus ABPP-BRET method provides the opportunities to study the function of the target protein with an unprecedented specificity in live cells in a longitudinal manner.

**Figure 1.**
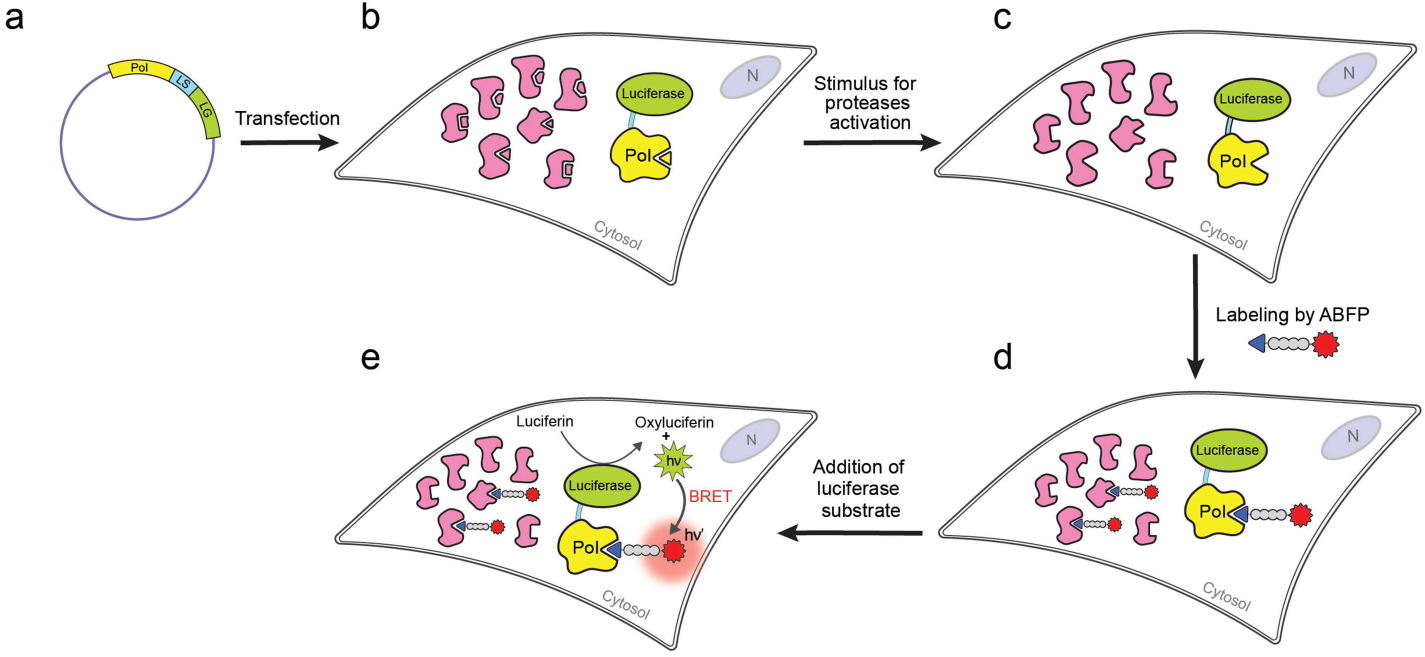
Schematic representation of ABPP-BRET technology. (a) Generation of a plasmid vector encoding for the luciferase tagged to the protease-of-interest (PoI); LS: linker sequence and LG: luciferase gene. Vector encoding the PoI tagged luciferase is expressed in a suitable cell line. (b) Cells expressing PoI tagged to luciferase enzyme along with the endogenous protease (pink) of the same or different families. Cells are subjected to an appropriate stimulus for the activation of PoI tagged to luciferase. (c) Cells expressing the active PoI tagged luciferase and other endogenous proteases. Cells are incubated with an activity-based fluorescent probe (ABFP) with an appropriate warhead and fluorophore. (d) Labeling of the PoI tagged luciferase enzyme along with other endogenous proteases of the same family by an ABFP. (e) Labeling of PoI by an ABFP creates an instantaneous BRET pair that reports the activity of the PoI.

## Results

### BRET Pair Selection

BRET assay is highly superior compared to FRET method because it has high signal to background ratio, as no external light is required; thus no auto fluorescence, photo bleaching of samples is not an issue and BRET assay can be used to study time kinetics or long term assay *in-vitro* and *in-vivo*^18^. Among the various factors, the choice of the donor (luciferase enzyme/substrate pair) and acceptor plays a significant role. We chose FMK-VAD-Rhodamine as an acceptor because this probe is commercially available for a reasonable price. It has been shown to target various cysteine proteases, including caspase (the target protein of our study).^16^ Most importantly, the rhodamine emission max is around 590 nm, highly suited for *in-vivo* BRET-based imaging applications. We chose RLuc 8.6 as a donor because this mutant luciferase has co-factor independent activity, high photon output, and red-shifted emission^19^. The RLuc8.6 uses coelenterazine h as a substrate, emitting at 535 nm^19^. Most importantly, the emission of coelentrazine h overlaps well with the absorption spectrum of rhodamine dye, an essential prerequisite for an efficient BRET process.

To obtain the RLuc8.6 emission spectrum, MCF-7 cells were transiently transfected with RLuc 8.6 plasmid. After 24 h of transfection, substrate coelenterazine h was added to the cells. Immediately after substrate addition, the emission from coelenterazine h was recorded. The emission maxima for the coelenterazine h were observed around 520-540 nm, consistent with the previous reports^19^. Afterwards, the absorption and emission spectra of rhodamine were acquired. We diluted the Rh-VAD-FMK probe in 1X PBS (Rh-VAD-FMK probe: 1X PBS, 1: 500 uL) and excited from 500-700 nm. The absorption maximum of rhodamine was observed at 561 nm, similar to the reported values. The coelenterazine h and rhodamine spectrum values were normalized and merged to evaluate the spectral overlapping between the coelenterazine h emission and the rhodamine excitation (Figure 2). The significant spectral overlap (540-600 nm) between the coelenterazine h and rhodamine establishes RLuc 8.6 and rhodamine as suitable BRET pairs.

**Figure 2.**
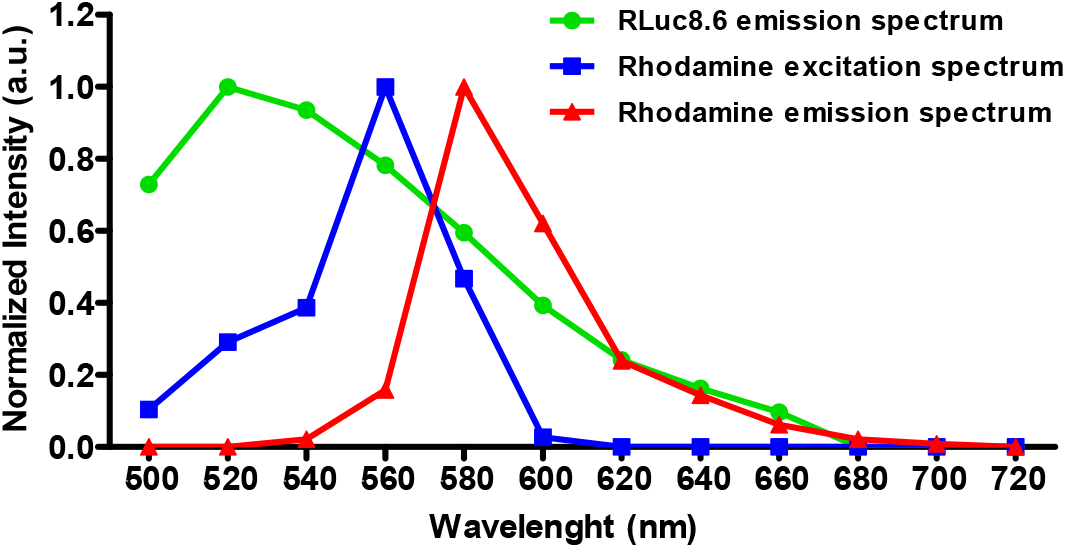
Spectral overlap between the normalized RLuc8.6 emission spectra and the normalized excitation and emission spectra of the rhodamine dye. The RLuc8.6 is excited by the addition of substrate, coelenterazine h, and the bioluminescence emission spectrum is recorded (green line). The rhodamine is excited with a 561 nm laser (blue line), and the emission spectra are recorded (red line). RLuc and rhodamine spectra are obtained separately using suitable filter combinations and merged to evaluate the spectral overlapping between the RLuc emission and rhodamine excitation and emission.

### Cloning and Expression of Caspase3-GGS-RLuc 8.6 Fusion Protein in MCF-7 Cells

To develop the BRET-ABPP, we chose caspase-3 as our protein of interest (PoI), as we have already demonstrated the caspase-3 activation via the FRET approach of ABPP^16,17^. Using suitable primers, the human caspase-3 gene was amplified by polymerase chain reaction (PCR) from pCMV3-C-OFPspark. The amplified caspase-3 gene was digested with restriction enzymes Hind III and Bgl II. The digested caspase-3 gene was cloned into the pCMV-GGS-RLuc 8.6 vectors between the Hind III and Bgl II restriction sites using PCR dependent restriction enzyme-based cloning method^20^. The resultant caspase3-GGS-RLuc 8.6 clone was confirmed by restriction digestion. We digested the suspected caspase3-GGS-RLuc8.6 plasmid with Hind III and Bgl II restriction enzymes as the caspase3 gene would be incorporated between these sites after ligation. After digestion, the caspase3 gene would be digested and give a fallout of around 850 bp (Figure 3a). We further confirmed the proper alignment and ligation of the caspase3 gene in the GGS-RLuc8.6 plasmid by carrying out DNA sequencing (supplementary figure 1 and supplementray data).

**Figure 3.**
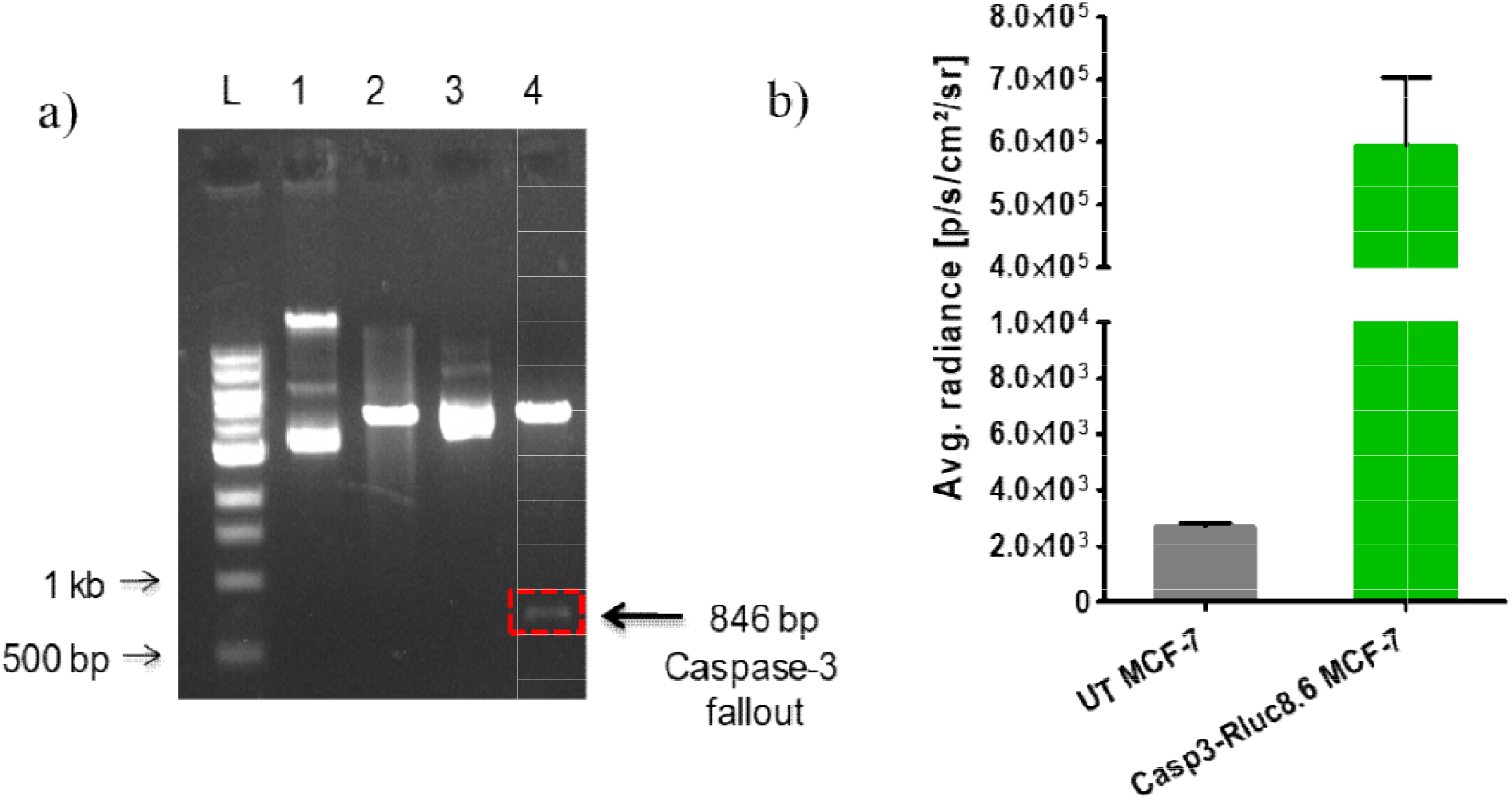
**a**. Recombinant pCMV-GGS-RLuc 8.6-caspase-3 plasmid was confirmed by restriction digestion analysis using Hind III and Bgl II Lane L, 1 kb DNA ladder; Lane 1, undigested pCMV-GGS-RLuc 8.6 plasmid; Lane 2, pCMV-GGS-RLuc 8.6 plasmid digested with Hind III and Bgl II; Lane 3, undigested recombinant plasmid pCMV-GGS-RLuc8.6-caspase-3; Lane 4, recombinant pCMV-GGS-RLuc8.6-caspase-3 plasmid digested with Hind III and Bgl II. **b**. The expression of recombinant RLuc8.6-GGS-caspase-3 fusion protein (Casp3-RLuc8.6) was determined by quantifying the bioluminescence. The bar graph representing the average radiance [p/s/cm^2^/sr] for untransfected (UT) MCF-7 cells and transfected MCF-7 cells.

The expression of the recombinant caspase-3-GGS-RLuc 8.6 fusion protein was determined by transiently transfecting MCF-7 cells with pCMV-caspase-3-GGSRLuc 8.6 plasmid. The transfected cells and control cells(untransfected) were seeded in 96-black well plate (20000 cells/well) and incubate for 24h at suitable conditions followed by the addition of coelenterazine h. Immediately after coelenterazine h addition, average radiance was recorded for both the untransfected and pCMV-caspase-3-GGS-RLuc 8.6 transfected MCF-7 cells. The average radiance values for the caspase-3-GGS-RLuc8.6 transfected cells (5.9 × 10^5^ photons/sec/cm^2^/sr) are significantly higher as compared to the untransfected cells (2.6 × 10^3^ photons/sec/cm^2^/sr), confirming that the expression of recombinant caspase-3-GGS-RLuc 8.6 protein in transfected MCF-7 cells (Figure 3 b).

### Monitoring of Real-Time Activation of Caspase-3 in Live Cells and Target Engagement Studies

To demonstrate the uniqueness of the ABPP-BRET method, we chose to study the activation of the caspase-3 enzyme in the MCF-7 cell line because it lacks an endogenous caspase-3 enzyme^21^. The transfected MCF-7 cells were treated with staurosporine (STS, pan-kinase inhibitor) (1 µM) for 4 h for the induction of apoptosis^22,23^. The STS triggers apoptosis via the intrinsic pathway and leads to the activation of caspases. Subsequently, cells were incubated with ABFP, Rh-VAD-FMK probe (BRET acceptor) for the labeling of PoI, and other proteases of the same family. The Rh-VAD-FMK probe was chosen because it has been previously shown that this probe labels caspase-3 with low selectivity^24,25^ and thus avoiding excessive engineering of ABFP probes. The RLuc 8.6 substrate coelenterazine h was added to the cells, and the average radiance was recorded in the donor emission channel and the acceptor emission channel. The BRET measurements were calculated using the mBRET values for the STS treated versus untreated samples^26^. The calculated mBRET value for the STS treated sample is 60 ± 0.4, which is significantly higher than the untreated control, 11 ± 9.2 (Figure 4a). This increase in mBRET values in the treated samples indirectly reports real-time activation of caspase-3 enzyme in live cells.

**Figure 4.**
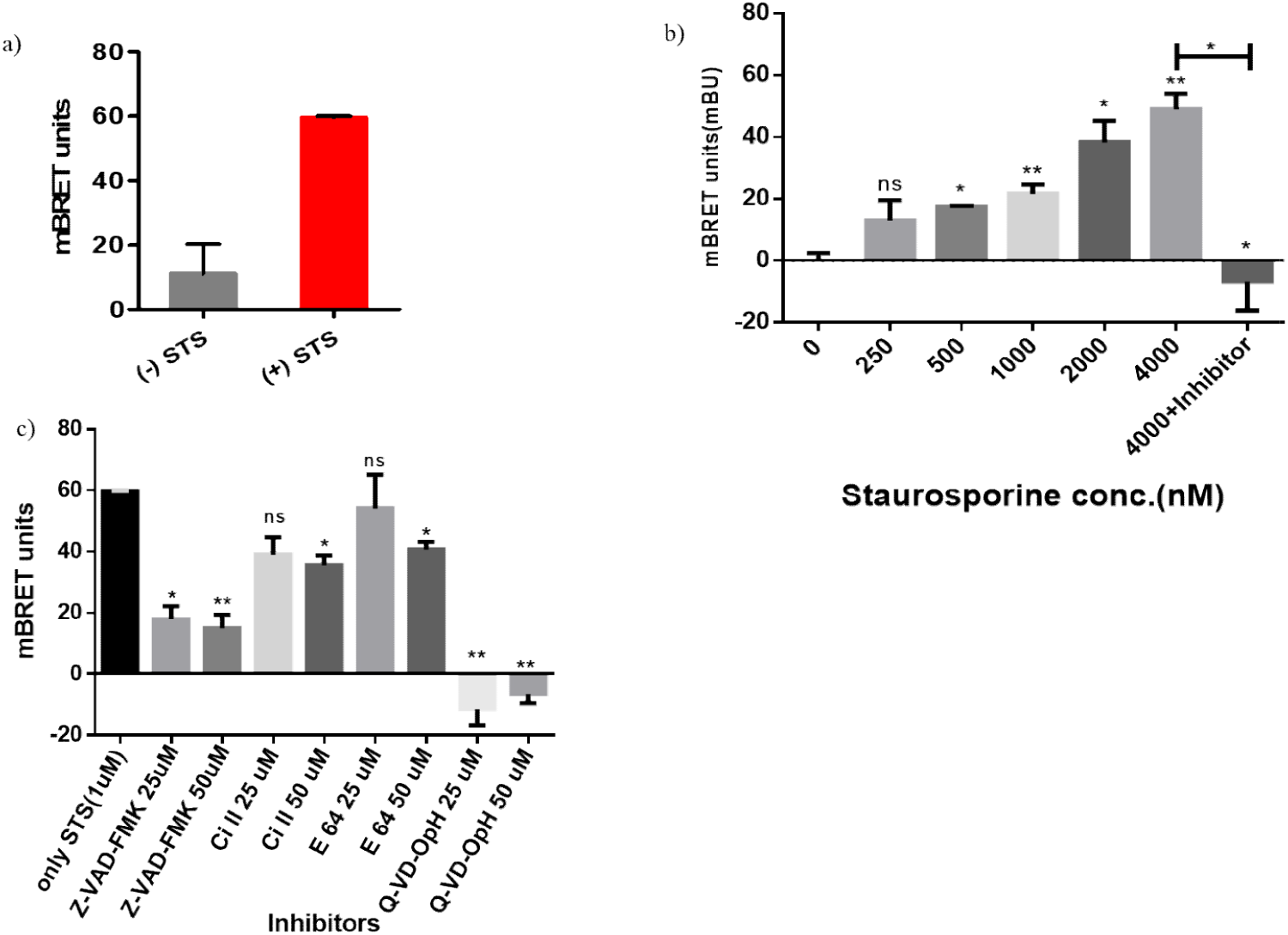
**a**. Caspase-3-GGS-RLuc8.6 transfected MCF-7 cells were treated with or without STS (1 µM) for 4 h. Subsequently, cells were incubated with the Rh-VAD-FMK probe for 1 h, followed by coelenterazine h addition. Immediately after coelenterazine h addition, average radiance was recorded, and the mBRET values for both the STS treated and untreated cells were calculated. **b**. Caspase-3GGS-RLuc 8.6 transfected MCF-7 cells were treated without and with different concentration of STS (250-4000 nM) for 4 h, and one group was treated with 4000 nM of STS for 4 h followed by addition of Z-VAD-FMK (1 mM) for 1 h. Subsequently, cells were incubated with the Rh-VAD-FMK probe for 1 h, followed by coelenterazine h addition. Immediately after coelenterazine h addition, average radiance was recorded, and the mBRET values for untreated cells and STS treated were calculated and a bar graph was generated **c**. Caspase-3 RLuc8.6 transfected MCF-7 cells treated with STS (4 µM) for 4 h. Cells were subsequently treated with different inhibitors, Z-VAD-FMK, caspase inhibitor I (Ci I), E-64 and Q-VD-OPh at two different concentrations 25 or 50 µM for 1 h. After inhibitor treatment, cells were incubated with the Rh-VAD-FMK probe for 1 h, followed by the substrate coelenterazine h addition. The average radiance was recorded in the acceptor and the donor channel, the mBRET values are calculated and represented as a bar graph.(ns-non significant, *-p value<0.05. ** -p value>0.01)

Overactivation of caspase-3 leads to several diseases such as acute neurological diseases such as Huntington’s disease^27^, Parkinson’s disease^28^, Alzheimer’s disease^29^, etc. The inhibitors that are capable of blocking caspase-3 activity would be potential drug candidates and hence beneficial for screening drug targets^30,31^. To check the effect of increasing concentration of STS on caspase activation, we treated MCF-7 cells expressing caspase-3-GGS-RLuc8.6 fusion protein with increasing concentration of STS for 4h. The transfected cells were treated with 250nM, 500nM, 1000 nM, 2000 nM and 4000 nM of STS; we treated one group of 4000 nM STS group with 1mM of Z-VAD-FMK (caspase inhibitor)^32^ for 1h and then treated with ABPP-BRET probe for 1h. BRET signal was recorded for vehicle control, treatment groups and inihibitor group. We observed dose-dependent increase in mBRET values in the treatmen group. However, the inhibitor group shown reduced BRET value (Figure 4b). This data suggests that the BRET-based approach can be applied to determine the activation as well as inhibition of caspase activity. Thus we can effectively use the ABPP-BRET method to screen caspase inhibitors and activators.

Developing a robust screening platform is extremely important for drug discovery programs. Many drugs fail during various phases of clinical trials due to a lack of efficacy. This failure is attributed to a lack of robustness in the screening platforms that can select true lead molecules. Therefore, developing a robust, highly sensitive screening method that can provide an absolute specificity is highly desired. Towards that goal, here we demonstrate the application of ABPP-BRET as a superb screening platform. For inhibitor screening, caspase-3-GGS-RLuc8.6 expressing MCF-7 cells were treated with STS (1 µM) for 4 h. Subsequently, after STS treatment, cells were incubated with Z-VAD-FMK or caspase inhibitor I or E64 or Q-VD-OPhH at two different concentrations (25 or 50 µM) for 1 h. These compounds were selected as they shown to inhibit caspase-3, other caspases and cysteine proteases.^32–35^ Cells were treated with the Rh-VAD-FMK probe for 1 h, followed by the addition of coelenterazine h. Immediately after coelentrazine h addition, the BRET signal was recorded.. In the absence of an inhibitor, the calculated mBRET value of the sample is 60 ± 0.4. The inhibition was seen at 50 *μ*M concentration by all inhibitors, but the maximum inhibition was shown by the Q-VD-OPh inhibitor, which was reported as a potent caspase3 inhibitor^36^. The mBRET values are lower for the Q-VD-OPh inhibitor as compared to the Z-VAD-FMK inhibitor, caspase inhibitor I and E-64 inhibitor (Q-VD-OPh > Z-VAD-FMK > caspase inhibitor I > E64 (Figure 4c). The above result state that the ABPP-BRET method can be used to measure quantitative inhibition of caspase-3, and this can be used to screen the inhibitor of caspases.

### Demonstration of ABPP-BRET in Stable Cells

The transient transfection method is a straightforward method for introducing a foreign gene onto mammalian cells. However, this method has several drawbacks, such as the transfected plasmid may be lost during replication and hence may not be suitable for longitudinal or *in-vivo* studies. In addition, over-expression of a certain gene may alter the physiology of the cells. Most importantly, this method cannot be used to generate helpful disease models in rodents. Considering these limitations, we thought demonstrating the ABPP-BRET method in stable cells is critical. The stable cells were generated by transfection of plasmid then selection of clones by selection marker^37^ as explained in the methods section. Three clones were selected, and their bioluminescence efficiency was measured. Clone 1 has shown high bioluminescence and is selected for further studies (Figure 5a). The MCF-7 cells stably expressing caspase3-GGS-RLuc8.6 (MCF-7 CR) were treated with PBS, and different concentrations of STS (50, 100, and 200 nM) of STS for 4 h and one group was treated with 200nM of STS followed by 1mM of Z-VAD-FMK (caspase inhibitor). Subsequently, cells were incubated with Rh-VAD-FMK probe for 1 h followed by the addition of coelenterazine h.. Immediately after substrate addition, the BRET signal was recorded, and the mBRET values were calculated and plotted as a bar graph (Figure 5b). We found an increase in mBRET values with increasing STS concentration. A ∼4-fold increase in mBRET was found at 200 nM of STS concentration compared to the untreated cells. The addition of a 1mM z-VAD-FMK inhibited the activation of caspase-3 and thus abolished BRET effect (figure 5b). The above results confirms the occurrence of BRET between RLuc 8.6-coelenterazine h and rhodamine and can be used to study activation or inhibition of caspase-3 activity in stable cells

**Figure 5.**
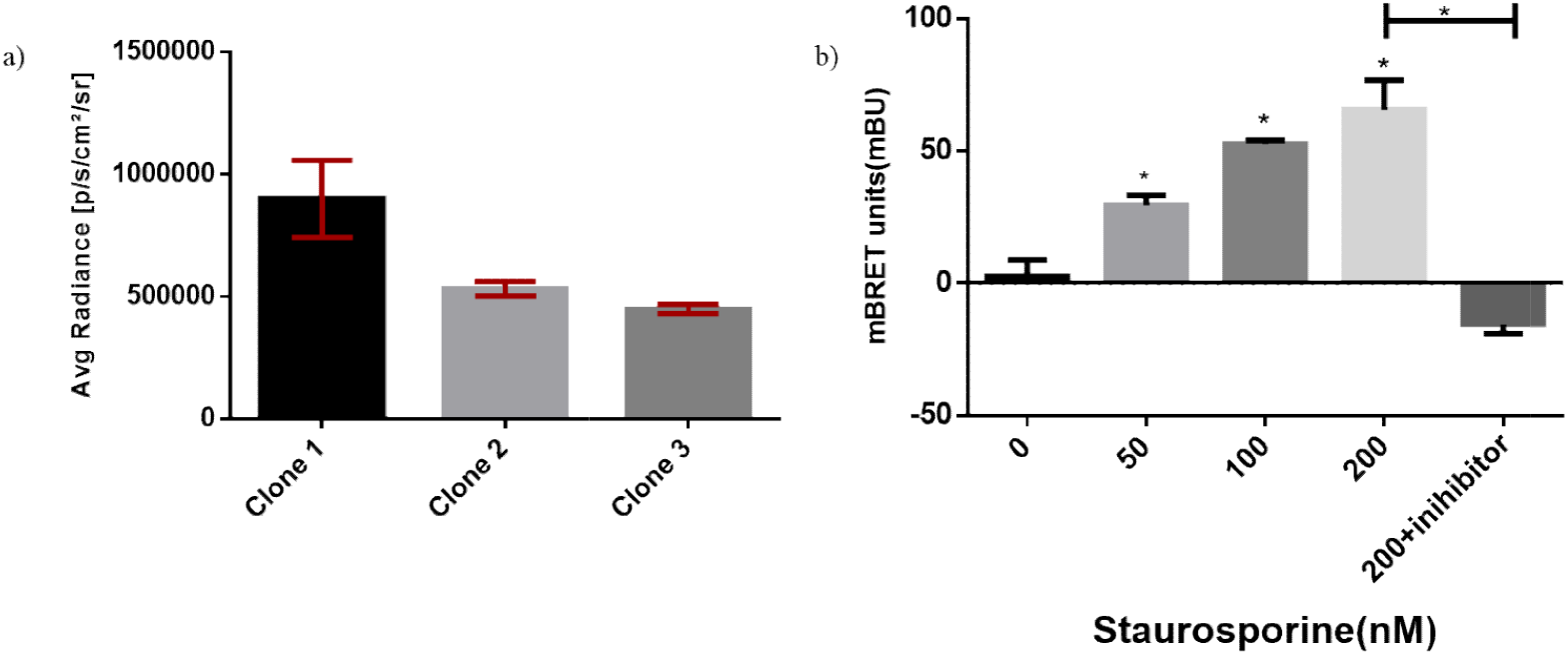
**a**. The 10,000 cells/well of MCF-7 Caspase3-GGS-RLuc8.6 clones, clone no. 1, 2 and 3 were seeded in 96 well plate and incubated for 16 h. After incubation, RLuc 8.6 substrate, coelenterazine h, was added in each well to measure luciferase activity. The average radiance for clone no. 1, 2 and 3 was recorded in the open filter and represented as a bar graph. **b**. 20000 cells of clone 1 were seeded in 96 well plate and were treated with increasing concentrations of STS (0, 50, 100, and 200 nM) for 4 h and one of the group was treated with 200 nM STS followed by addition of 1 mM Z-VAD-FMK inhibitor. Subsequently, cells were incubated with the Rh-VAD-FMK probe for 1 h, followed by coelenterazine h addition. Immediately after coelenterazine h addition, average radiance was recorded, and the mBRET values were calculated and represented as a bar graph. (ns-non significant, * p value < 0.05)

## Discussion

Even after two decades of the complete human genome draft, we still do not know the function of many proteins. The genomics^38,39^, transcriptomics, and standard proteomics technologies^40,41^ have failed to provide information on the function of many proteins. This is mainly attributed to the reason that the function of “active protein’ is regulated at multiple levels such as irreversible post-translational modification^42^, reversible interaction of active proteins with endogenous inhibitors^43,44^. ABPP-MS method addresses these limitations posed by major “omics” technologies and helped to decipher the function of many proteins, which are involved in many human diseases^5^. ABPP-MS has succeeded as a robust mature technology over the last decade. Because of its unparalleled strength, many academic laboratories and pharmaceutical/biotech industries routinely use ABPP-MS for target identification, target validation, and inhibitor screening^6–12^. However, the existing ABPP-MS method uses fractionated proteomes from cells/tissues for downstream analysis. Because of this shortcoming, cells and tissues are subjected to sample homogenization and this leads to a loss of spatial information on protein activity^45,46^. In addition, the current ABPP-MS method cannot be used to detect the function of protein directly from living systems in real-time in a longitudinal manner.

Considering these limitations it is important to develop next-generation activity-based chemical proteomics technology, which does not rely on gel- or MS-based detection platforms. Towards that goal, many research groups are developing complementary technologies. Our group had developed a different version of the activity-based chemical proteomics method named AbRGT 1.0.^16,17^ which utilizes FRET-based read out to detect protein activity. Although powerful, the AbRGT-FRET method has certain limitations and therefore reduces the scope of this technology. Therefore, the present study was directed towards preserving superior attributes of activity-based reporter gene technology (AbRGT) while addressing some of the limitations caused by FRET-based measurement and avoiding the use of a highly expensive confocal microscope. To do that, we decided to explore bioluminescence resonance energy transfer (BRET) technology. The BRET method is currently used for the detection of non-invasive imaging of protein function both in live cells as well as animal or plant models in real-time in a longitudinal manner^48,49^. However, most of the previous BRET-based method lack specificity and therefore produces ambiguous results. The pioneering work by Robers and coworkers address the “specificity issue” by developing nanoBRET platforms.^50-52^ This method provides unprecedented specificity to study the function of active proteins and can be used for target engagement studies and inhibitor profiling in live cells in real-time. However, one of the major limitations of this approach is the lengthy and inflexible synthesis of chemical probes against each enzyme, which makes this method not very appealing^50^. In this regard, we would like to develop a technology, which can be applied to a large number of proteins by using commercially available, off-the-shelf family-wide fluorescent activity-based probes. To achieve this goal, we decided to combine ABPP with the BRET method to create ABPP-BRET technology. As a proof concept, we transiently expressed caspase-3-luciferase enzyme in the MCF-7 cells followed by treatment with a pan-kinase inhibitor (STS). The treatment of MCF-7 cells with STS leads to proteolytic activation of caspase-3-luciferase enzyme followed by labeling of caspase-3 enzyme of the fusion protein by FMK-VAD-Rhodamine chemical probe to create an in-situ BRET pair. As expected, there was no or negligible BRET before probe addition however we observed an increase in BRET value with an increased concentration of STS. To demonstrate that the BRET signal truly comes from the target protein (i.e. caspase-3-luciferase enzyme), we have carried out experiments in the presence of potent caspase inhibitors, as expected, the BRET signal considerably decreased in the presence of various caspase inhibitors. This result manifests that the BRET signal is indeed coming from the target protein. Further, we have utilized this method for target-engagement studies with various irreversible and reversible inhibitors. As expected, we obtained different mBRET values depending on the compound’s potency. Further, we created an engineered stable MCF-7 cell line expressing caspase-3 luciferase fusion protein followed by demonstration of dose-dependent proteolytic activation of caspase-3-luciferase enzyme upon treatment with different concentrations of STS.

It is pretty evident that these three technologies (ABPP-MS, ADPL, and ABPP-BRET) have their own merits and limitations. For example, ADPL^13–15^ and ABPP-BRET cannot be used for target identification in an unbiased manner at the system level whereas ABPP-MS^6–12^ is routinely used for this purpose. Similarly, ABPP-MS and BRET-ABPP cannot be employed for the analysis of patients’ samples whereas ADPL is uniquely suited to do that^15^. ABPP-BRET can be used for profiling proteins in live cells in real-time in a longitudinal manner but the other two technologies do not have that capability. ABPP-MS and ADPL do not involve the genetic engineering of cells and therefore provide the opportunity to look at the activity of endogenous proteins whereas ABPP-BRET involves the expression of exogenous fusion reporter protein and just provides a proxy for endogenous protein activity. Therefore ABPP-BRET technology is not a replacement for the most powerful ABPP-MS or ADPL technologies whereas it is complementary to the existing technologies. We firmly believe each of these three technologies complements each other and therefore collectively provides powerful tools to interrogate protein function at the system level.

## Conclusion

In summary, we have introduced a new chemical proteomics technology to the existing toolbox. The ABPP-BRET technology provides an opportunity to study the function of “active enzyme” with unprecedented specificity. In particular, this method allows real-time monitoring of proteolytic activation of target protein in live cells. We also demonstrated that this method could be used for target engagement studies in live cells in real-time. Further, the creation of a stable cell line expressing caspase-3 luciferase offers longitudinal imaging of live cells and opportunities to translate the ABPP technology to follow the function of “active proteins” in live animals in real-time in a longitudinal manner with an unprecedented specificity.

## Materials and methods

### Reagents

Lipofectamine 2000, E-64 protease inhibitor, Q-VD-OPh hydrate, cpm-VAD-CHO, staurosporine and caspase-3 inhibitor I, coelentrazine h are procured from Sigma. Rh-VAD-FMK probe and Z-VAD-FMK inhibitor was obtained from Abcam.

### Cloning

For pCMV-caspase-3-GGS-RLuc 8.6 recombinant vector preparation, 846 bp of the human caspase-3 gene was PCR amplified from pCMV3-C-OFPspark plasmid using suitable primers. The amplified 846 bp caspase-3 PCR product was digested with Hind III and Bgl II restriction enzymes and cloned into pCMV-GGS-RLuc 8.6 vector between the restriction sites Hind III and Bgl II using a PCR based method for restriction cloning^20^. The clones are confirmed by restriction digestion and DNA sequencing.

### Transient transfections

MCF-7 cells were seeded in a 35 mm dish and were transiently transfected with pCMV-caspase-3-GGS-RLuc 8.6 recombinant plasmid at 70% cell confluency using Lipofectamine 2000 reagent. 24 h post-transfection, the cells were trypsinized and distributed in a 96 well black plate with a clear bottom (20,000 cells/well). Cells were allowed to attach by overnight incubation in a humidified atmosphere of 5% CO_2_ at 37 ºC before the start of experiments.

### Luciferase assay

In transfected MCF7 seeded in 96-black well plate, RLuc 8.6 substrate, coelenterazine h (50 µL of the 1 mg/mL stock, dissolved in 1X PBS) was added to each well. Immediately after substrate addition, cells were analyzed in an IVIS spectrum with an emission spectrum (520-540 nm) using an integration time (60 sec/filter) and and the average radiance for each well was recorded.

### Caspase-3 activity measurement

For the caspase-3 activity measurements, pCMVcaspase-3-GGS-RLuc 8.6 transfected MCF-7 cells in a 96 black well clear bottom plate were treated with STS (1 µM) for 4 h. Cells were then incubated with the Rh-VAD-FMK probe for an additional 1 h. Subsequently, RLuc 8.6 substrate, coelenterazine h, was added to the cells, and the average radiance value for each well was taken immediately using an IVIS spectrum at 520-540 nm wavelength as donor filter and 560-580nm as acceptor filter.

### Effect of inhibitor on STS based caspase activation

The MCF7 cells transfected with pCMV-caspase3-GGS-Rluc8.6 plasmid were seeded in 96 black well plate clear bottom(20000 cells/well) and treated with PBS or increasing concentration of STS (250-4000 nM) for 4 h. One group of treated cells were incubated for 1 h with Z-VAD-FMK(50μM). Both the groups were incubated with ABFP, Rh-VAD-FMK probe for 1 h. The RLuc 8.6 substrate coelenterazine h was added to the cells, and the average radiance was recorded in the donor emission channel, RLuc 8.6 (520-540 nm), and the acceptor emission channel, rhodamine (580-600 nm) channel. BRET calculations were performed as described before.

### Inhibitors screening

For inhibitors screening, pCMV-caspase-3-GGS-RLuc 8.6 transfected MCF-7 cells in 96 black well plates with clear bottom were pretreated with different inhibitors. The treatment of the inhibitors was given for 1 h before Rh-VAD-FMK probe incubation. Different concentrations of inhibitors (25 and 50 µM) were tested. Subsequently, the caspase-3 labeling was performed by incubation of the cells with the Rh-VAD-FMK probe for 1 h. For BRET studies, coelentrazine h was added to the cells, and immediately after substrate addition, the average radiance was recorded.

### Preparation of stable clone

The MCF7 cells were seeded in a 35 mm plate at 70% confluency and subsequently transfected with pCMV-caspase3-GGS-RLuc8.6 plasmid by using Lipofectamine 2000 as per manufacturer’s instructions. The transfected cells were incubated for 48 h for the expression of the transfected plasmid. Transfected cells were then grown in RPMI under zeocin (300 µg/ml), an antibiotic selection marker, to allow colony formation from single cells in a 10 cm plate. The plate was incubated till colonies were formed. Colonies formed from single cells were picked up and grown in 96-well plate. After cells attain confluency in a 96-well plate, they were transferred to a 24-well plate, and then clones were grown in bulk to check the expression of caspase-3-GGS--Rluc8.6 protein with respect to luciferase activity of RLuc8.6. The clones with high luciferase activity were selected for further experiments.

### BRET measurements

To perform the BRET measurements for each sample, the image obtained from IVIS spectrum was analysed using Living Image software (version 4.5). The image file was opened and region of interest (ROI) were drawn over each well, and the values of average radiance (photons/sec/cm^2^/sr) were obtained. The average radiance values were acquired for the donor channel (donor emission channel, 520-540 nm, RLuc8.6), and the acceptor channel (acceptor emission channel, 580-600 nm, rhodamine). The acceptor/donor channel emission (A/D) ratio was calculated for the donor (D) only, and the donor + acceptor (D+A) sample. A/D channel emission ratio of the D only sample was subtracted from the D+A sample to obtain the corrected BRET ratio (cBRET)^26^. Subsequently, mBRET values were calculated by multiplying the values of the corrected BRET ratio to 1000. The calculations were done using MS excel; data is statistically analyzed and represented using Graph Prism software version 6.5.

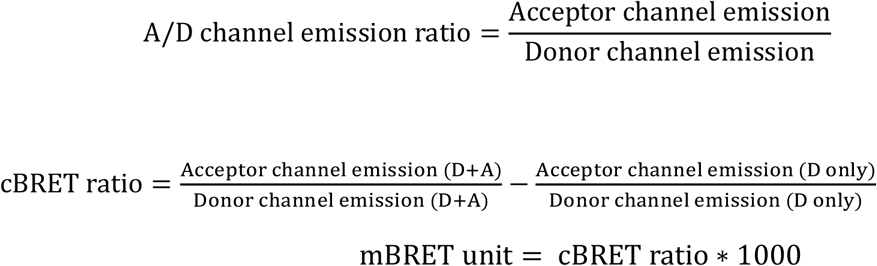

## Acknowledgments

This work was supported DST-SERB Early Career Award to B.S.S., DST-SERB (ECR/2015/000253), and the DBT grant (BT/PRI1450/BRB/I0/1370/2015).

