## Supplementary material for "Development of Non-invasive ABPP-BRET Platform Technology Enable Real-Time Detection of Post-Translational Modification and Target Engagement with Absolute Specificity in Living Systems": Supp Info

Table of Contents page no


1. Table 1 Primers used for Caspase3 cloning page 2
2. Table 2 PCR thermocycler conditions for amplification of Caspase3 gene 2
3. Figure 1 Gel image of PCR amplification of Caspase3 3
4. Figure 2 Gel image of restriction digestion of pCMV-GGS-RLuc8.6 3
5. Figure 3 Gel of colony PCR done of transformants from ligation 4
6. Figure 4 Gel image of restriction digestion of selected positive colonies 5
7. Figure 5 Nucleotide blast result 6
8. Figure 6 Plasmid map of pCMV-Caspase3-GGS-RLuc8.6 7
9. Table 3 Information of caspase inhibitors used in the study 7
10. Figure7 Structure of caspase inhibitors used in study 8

Supplementary Table 1 - Sequence of primers used for cloning

| Primer name | DNA sequence |
|---|---|
| FCasp3HindIII | 5GCCAAGCTTGCCACCATGGAGAACACTGAAAACTC3 |
| RevCasp3BglII | 5GCCAGATCTGTGATAAAAATAGAGTTCTTTTGTG3 |

**Supplementary Table 1** - Sequence of primers used for cloning

| PCR thermocycler conditions | | | |
|---|---|---|---|
| | **Steps** | **Temp.** | **Time** |
| 1 | Denaturation<br>30 cycles | 95 C | 4 mins |
| 2 | Denaturation | 95 C | 45 secs |
| 3 | Annealing | 60° C to 70°C | 15 secs |
| 4 | Elongation<br>30 cycles | 72 | 30 secs |
| 5 | Elongation | 72 | 2 mins |
| 6 | Storage | 4 | ∞ |

**Supplementary Table 2** - Table for standardization of PCR thermocycler conditions for amplification of caspase-3 gene.

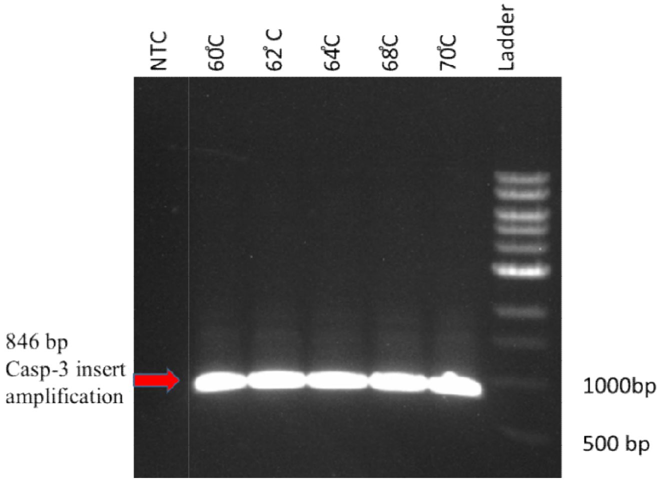


**Supplementary Figure 1-** Agarose gel image of PCR amplicon from different annealing temperature.

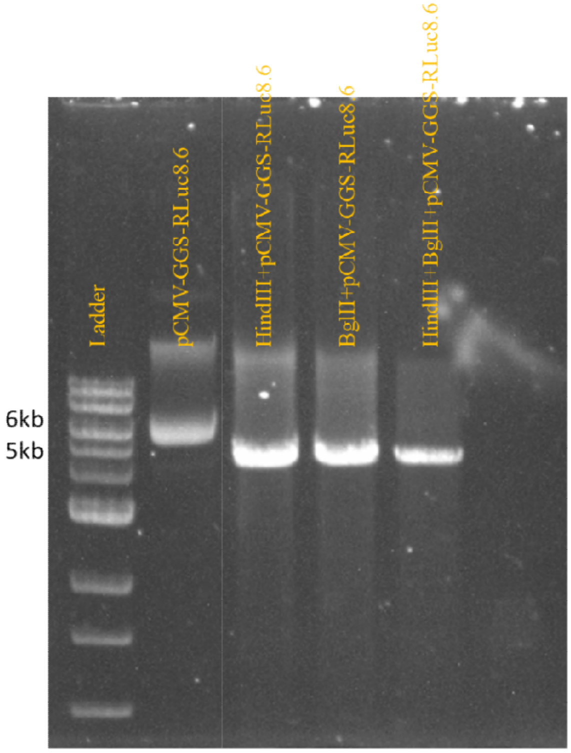


**Supplementary Figure 2 -** Agarose gel electrophoresis image of pCMV-GGS-RLuc8.6 plasmid digested with HindIII and BglII restriction enzyme.

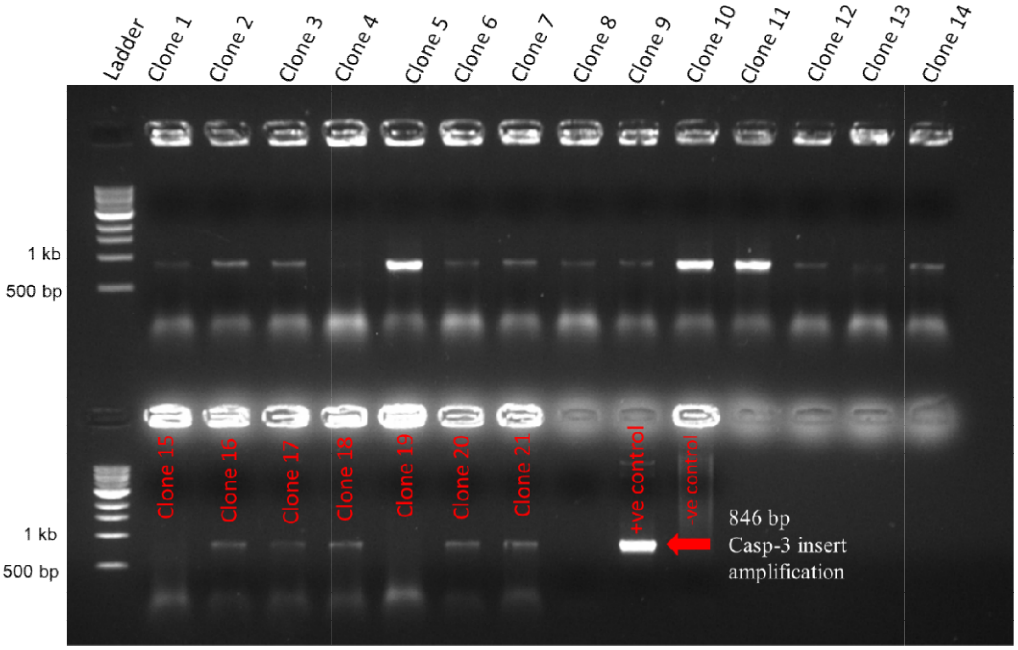


**Supplementary Figure 3** - Agarose gel electrophoresis image of colony PCR done of transformants from ligation of pCMV-GGS-RLuc8.6 and caspase 3 PCR amplified insert.

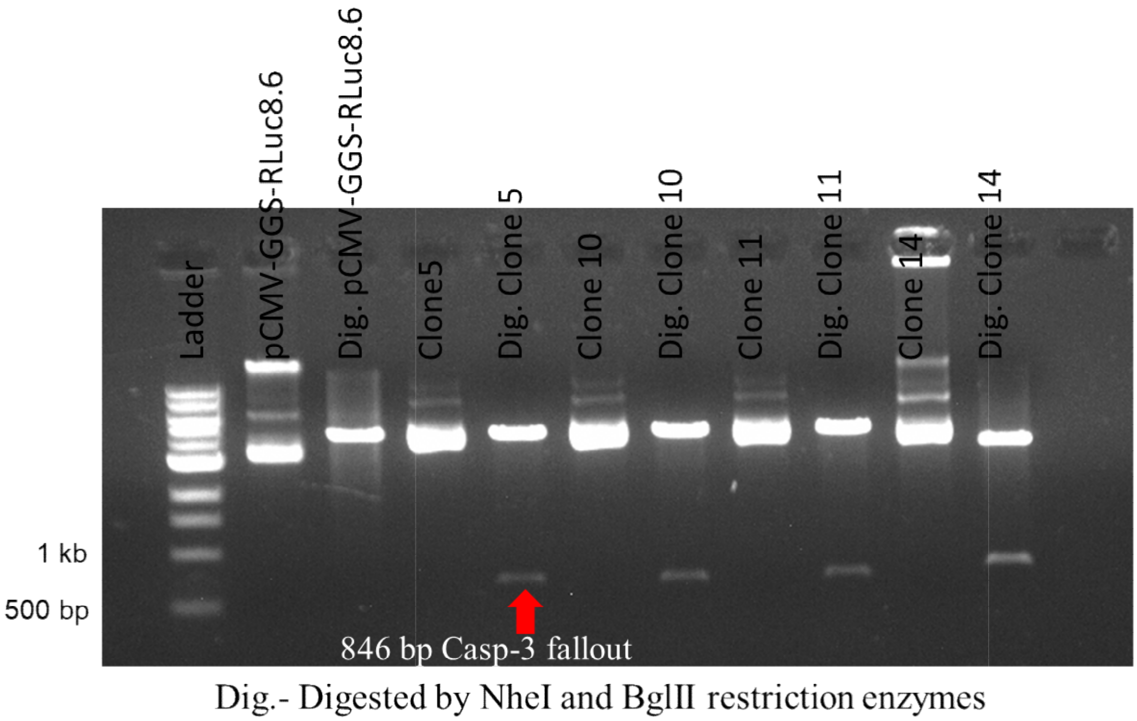


**Supplementary Figure 4** - Agarose gel electrophoresis image of digestion of selected positive colonies from Colony PCR with NheI and BglII restriction enzyme.

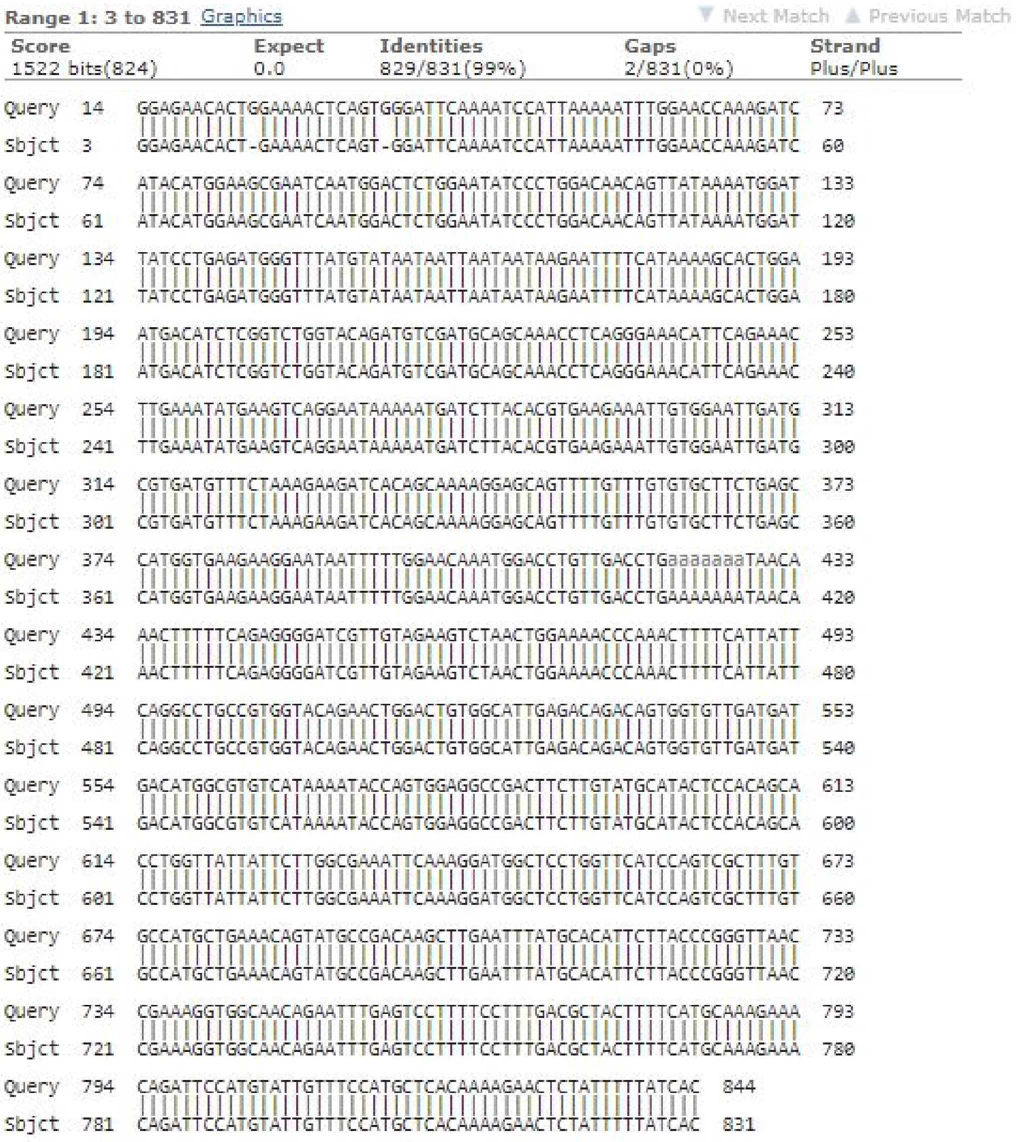

```
Range 1: 3 to 831 Graphics                              Next Match  Previous Match
Score            Expect   Identities       Gaps          Strand
1522 bits(824)   0.0      829/831(99%)     2/831(0%)     Plus/Plus

Query  14   GGAGAACACTGGAAAACTCAGTGGGATTCAAAATCCATTAAAAATTTGGAACCAAAGATC  73
            |||||||||| ||||||||||| |||||||||||||||||||||||||||||||||||||
Sbjct  3    GGAGAACACT-GAAAACTCAGT-GGATTCAAAATCCATTAAAAATTTGGAACCAAAGATC  60

Query  74   ATACATGGAAGCGAATCAATGGACTCTGGAATATCCCTGGACAACAGTTATAAAATGGAT  133
            ||||||||||||||||||||||||||||||||||||||||||||||||||||||||||||
Sbjct  61   ATACATGGAAGCGAATCAATGGACTCTGGAATATCCCTGGACAACAGTTATAAAATGGAT  120

Query  134  TATCCTGAGATGGGTTTATGTATAATAATTAATAATAAGAATTTTCATAAAAGCACTGGA  193
            ||||||||||||||||||||||||||||||||||||||||||||||||||||||||||||
Sbjct  121  TATCCTGAGATGGGTTTATGTATAATAATTAATAATAAGAATTTTCATAAAAGCACTGGA  180

Query  194  ATGACATCTCGGTCTGGTACAGATGTCGATGCAGCAAACCTCAGGGAAACATTCAGAAAC  253
            ||||||||||||||||||||||||||||||||||||||||||||||||||||||||||||
Sbjct  181  ATGACATCTCGGTCTGGTACAGATGTCGATGCAGCAAACCTCAGGGAAACATTCAGAAAC  240

Query  254  TTGAAATATGAAGTCAGGAATAAAAATGATCTTACACGTGAAGAAATTGTGGAATTGATG  313
            ||||||||||||||||||||||||||||||||||||||||||||||||||||||||||||
Sbjct  241  TTGAAATATGAAGTCAGGAATAAAAATGATCTTACACGTGAAGAAATTGTGGAATTGATG  300

Query  314  CGTGATGTTTCTAAAGAAGATCACAGCAAAAGGAGCAGTTTTGTTTGTGTGCTTCTGAGC  373
            ||||||||||||||||||||||||||||||||||||||||||||||||||||||||||||
Sbjct  301  CGTGATGTTTCTAAAGAAGATCACAGCAAAAGGAGCAGTTTTGTTTGTGTGCTTCTGAGC  360

Query  374  CATGGTGAAGAAGGAATAATTTTTGGAACAAATGGACCTGTTGACCTGaaaaaaaTAACA  433
            ||||||||||||||||||||||||||||||||||||||||||||||||||||||||||||
Sbjct  361  CATGGTGAAGAAGGAATAATTTTTGGAACAAATGGACCTGTTGACCTGAAAAAAATAACA  420

Query  434  AACTTTTTCAGAGGGGATCGTTGTAGAAGTCTAACTGGAAAACCCAAACTTTTCATTATT  493
            ||||||||||||||||||||||||||||||||||||||||||||||||||||||||||||
Sbjct  421  AACTTTTTCAGAGGGGATCGTTGTAGAAGTCTAACTGGAAAACCCAAACTTTTCATTATT  480

Query  494  CAGGCCTGCCGTGGTACAGAACTGGACTGTGGCATTGAGACAGACAGTGGTGTTGATGAT  553
            ||||||||||||||||||||||||||||||||||||||||||||||||||||||||||||
Sbjct  481  CAGGCCTGCCGTGGTACAGAACTGGACTGTGGCATTGAGACAGACAGTGGTGTTGATGAT  540

Query  554  GACATGGCGTGTCATAAAATACCAGTGGAGGCCGACTTCTTGTATGCATACTCCACAGCA  613
            ||||||||||||||||||||||||||||||||||||||||||||||||||||||||||||
Sbjct  541  GACATGGCGTGTCATAAAATACCAGTGGAGGCCGACTTCTTGTATGCATACTCCACAGCA  600

Query  614  CCTGGTTATTATTCTTGGCGAAATTCAAAGGATGGCTCCTGGTTCATCCAGTCGCTTTGT  673
            ||||||||||||||||||||||||||||||||||||||||||||||||||||||||||||
Sbjct  601  CCTGGTTATTATTCTTGGCGAAATTCAAAGGATGGCTCCTGGTTCATCCAGTCGCTTTGT  660

Query  674  GCCATGCTGAAACAGTATGCCGACAAGCTTGAATTTATGCACATTCTTACCCGGGTTAAC  733
            ||||||||||||||||||||||||||||||||||||||||||||||||||||||||||||
Sbjct  661  GCCATGCTGAAACAGTATGCCGACAAGCTTGAATTTATGCACATTCTTACCCGGGTTAAC  720

Query  734  CGAAAGGTGGCAACAGAATTTGAGTCCTTTTCCTTTGACGCTACTTTTCATGCAAAGAAA  793
            ||||||||||||||||||||||||||||||||||||||||||||||||||||||||||||
Sbjct  721  CGAAAGGTGGCAACAGAATTTGAGTCCTTTTCCTTTGACGCTACTTTTCATGCAAAGAAA  780

Query  794  CAGATTCCATGTATTGTTTCCATGCTCACAAAAGAACTCTATTTTTATCAC  844
            |||||||||||||||||||||||||||||||||||||||||||||||||||
Sbjct  781  CAGATTCCATGTATTGTTTCCATGCTCACAAAAGAACTCTATTTTTATCAC  831
```

**Supplementary Figure 5 -** Nucleotide blast result of plasmid caspase3-GGS-RLuc8.6 (subject) with caspase3 gene (query)

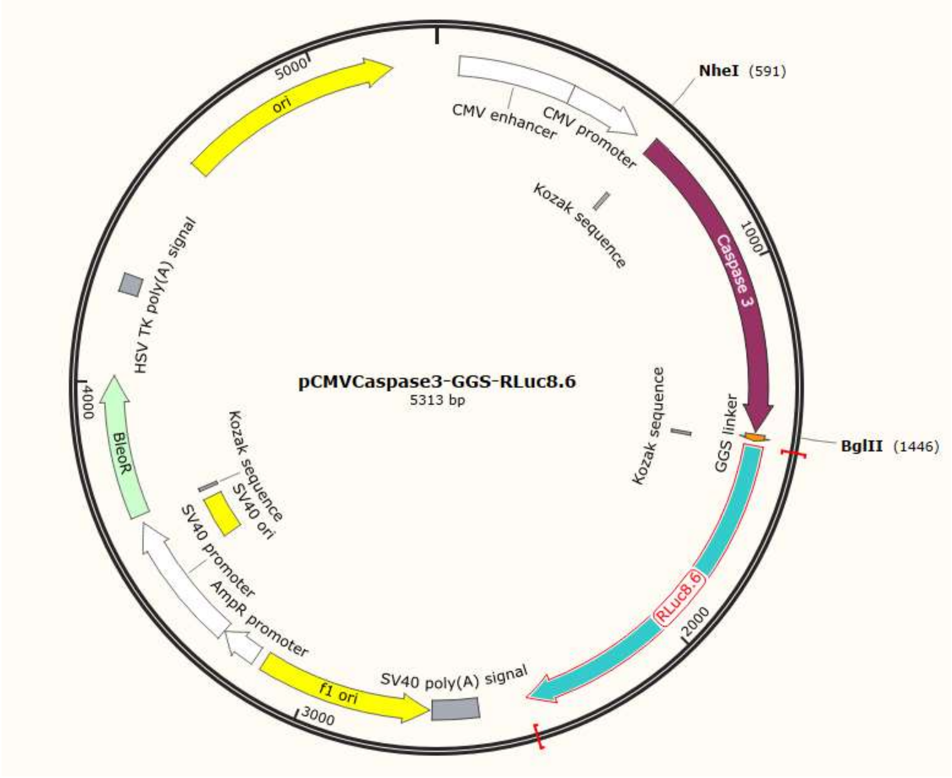


**Supplementary Figure 6 -** Plasmid map of pCMV-Caspase3-GGS-RLuc8.6

Supplementary table 3 Information of caspase inhibitors used in the study

| S.No. | Inhibitor type | Catalog no. | Reversible/Ireversible |
|---|---|---|---|
| 1 | E-64 protease inhibitor | E3132-1MG | Irreversible |
| 3 | Q-VD-OPh hydrate | SML0063-1MG | Irreversible |
| 4 | Caspase-3 Inhibitor I - CAS 169332-60-9 - Calbiochem | 235420-1MG | Reversible |

**Supplementary Table 3 -** Information of caspase inhibitors used in the study

E-64 protease inhibitor

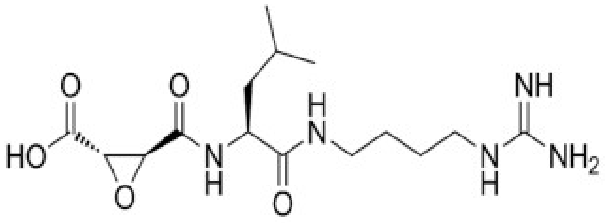


Q-VD-OpHydrate

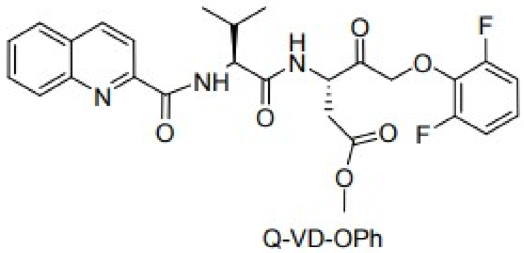


Caspase 3 inhibitor I

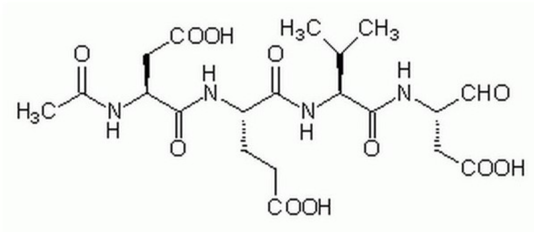


**Supplementary Figure 7** - Structure of caspase inhibitors used in study
